# Reactivation of atrium genes is a primer for heart infarction or regeneration

**DOI:** 10.1101/2020.04.02.021030

**Authors:** Yisong Zhen

## Abstract

The inability of the adult heart to repair or regenerate is manifested in prevalent morbidity and mortality related to myocardial infarction and heart failure. However, the cue to the reactivation of cardiomyocyte proliferation in the adult remains largely unknown. In the present study, three independent datasets were explored using bioinformatics analysis methods to solve the problem. Our results revealed that atrium genes were upregulated in response to the injury, which indicates the possible cell type withdraw and reinitiation of proliferation capability. Our findings might provide an alternative viewpoint on the cardiomyocyte regeneration or myocardial infarction.

## Introduction

Cardiovascular diseases have long been the leading cause of death in both industrialized and developing countries. Underlying basis of the diseases is cardiomyocyte loss. The notion that it might be possible to rebuild the cardiac muscle paves the way of new concept, heart regeneration, which is one of the most active and controversial areas of biomedical research. Physician scientists at the bench-side propose three strategies, including cell-based therapy^1,2^, bioengineering methods^3,4^ and synthetic organogenesis^5–7^.

However, a major barrier against myocardial regeneration appears to be cell-cycle arrest of adult mammalian cardiomyocytes. One theory is that mitosis of incessantly contractile cells is infeasible^8^. Another theory is that neonatal heart regeneration in mammals is simply a remnant of developmental programs that were lost after birth^9,10^. Researchers around the world now verify them by using a wide variety of animal models^11^.

In zebrafish model, differentiated atrial cardiomyocytes can transdifferentiate into ventricular cardiomyocytes to contribute to cardiac ventricular regeneration^12^. Another group found that a subpopulation of cardiomyocytes that transiently express atrial myosin heavy chain (*amhc*) contributes to specific regions of the ventricle^13^. These finding suggests an unappreciated level of plasticity during chamber formation, and we therefore hypothesize that there might be a kind of heart atrium progenitor cell exists in the mammalian system. The progenitor cell will hence play a role in heart regeneration in response to injury stimuli. We designed the following study using public data to explore the possibility and revealed that heart regeneration came into being along with the emergence of atriumizaiton, upregulation of atrium specific genes.

## Methods

### Data sources

All three datasets used in the present study were downloaded from Gene Expression Omnibus (GEO). The dataset GSE1479 is a benchmark set for early cardiac development. At stage 10.5, the rostral and caudal parts of the mouse embryo were removed and the middle part, which includes the heart, was subjected to expression analysis. From embryonic day 11.5 on, embryonic hearts were isolated and the ventricular from the atrial chambers were separated. The GSE775 dataset is a time series experiment design intended to compare normal functioning left ventricles(lv) with infarcted (ilv) and non-infarcted left ventricles(nilv). Ilv samples are taken from the region between the LAD artery and the apex on a mouse with myocardial infarction. Nilv samples are taken from the region above the infarction and the left ventricle(lv) samples mimic that region in a sham mouse. Both GSE1479 and GSE775 were released by the CardioGenomics project^14^. The GSE64403 was published by Boyer’s lab on the subject of transcriptional reversion of cardiac myocyte fate during mammalian cardiac regeneration^15^. Primary myocardial tissues sampled from neonatal mice and murine hearts undergoing post-injury regeneration were harvested according to the original description.

#### Collation of atrium specific genes

Atrium (specific) genes are highly expressed in heart atrium compared with these in the ventricle. Therefore, atrium genes play import roles in chamber physiological function or morphogenesis during heart development. The GSE1479 is a benchmark set for early cardiac development. From embryonic day 11.5 on, embryonic hearts were isolated and separated to the ventricular from the atrial chambers. In the bioinformatic curation procedure, the microarray data from respective chambers were pooled together to perform gene differential expression analysis using limma package^16^. The expression value contrast above 1 (logFC > 1) and *p*-value above 0.001 were chosen as the complete chamber specific genes. After filtering, the hierarchical clustering was used to separate the atrium and ventricle sets. Manual curation of chamber specific genes was undertaken to construct an independent evaluation set.

### Source code management

The source code of the present project was deposited in the GitHub (atriumization_source_code.R^17^). The development process tried to follow the etiquette of the scientific reproducible research^18^.

### Data analysis tools

Data was analysis using the R/Bioconductor packages. Picture and graph layout were generated by ggplot2 and cowplot packages. Differential gene expression analysis was conducted using limma if the data type is Affymetrix microarray, or DESeq2 if the data is high-throughput sequencing. Gene Set Enrichment Analysis (GSEA) is a computational method that determines whether an *a priori* defined set of genes shows statistically significant, concordant differences between two biological states. This approach was conducted using ClusterProfiler package. Heatmap analysis was generated using the pheatmap package. Principle component analysis results were generated using the factoextra and FactoMineR packages. Most packages were downloaded from Bioconductor or CRAN^19^.

## Results

### Atrium specific genes curated from the public dataset

Principle component analysis (PCA) was performed to check the data quality (GSE1479). The original experiment claimed to monitor changes in gene expression related to heart development and maturation. From embryonic day 11.5 on, the expression profiling by array also reported the gene expression respectively from ventricular and the atrial chambers. In the present PCA results, components PC1 and PC2 can perfectly separate these two events, cardiac maturation and chamber differentiation although the maturation as an independent variable was partially collineared with chamber differentiation (Figure 1A). To minimize the co-founder of the maturation event, the chamber specific samples at different developmental stages were pooled together and gene differential expression analysis was performed to curate the chamber specific genes and then hierarchical clustering was conducted to obtain these two groups. After above filtering, the atrium and ventricle specific gene sets were generated accordingly. Atrium gene set included 189 genes with unique Entrez gene ID. Ventricle gene set included 99 genes, and the ratio of atrium and ventricle gene number is about 2:1 (Table S1). The chamber specific set was then used in the next round of PCA analysis. This resulted in the increase percentage of PC1 component and about 70 % variability can be explained by the new PC1. In the atrium gene set, *sarcolipin* (*Sln*, gene ID: 66402) is included, which is a hallmark of heart atrium^20,21^. To further confirm these findings, panorama view by the DAVID functional analysis was performed (Figure 1C & Figure 1D). GO term ‘cardiac chamber morphogenesis’ and ‘heart morphogenesis’ were enriched in the result. The term ‘secretion’ was enriched in protein function using DAVID analysis, which was in consistent with the previous observation that atrium is a secretory organ^22^.

**Figure 1.**
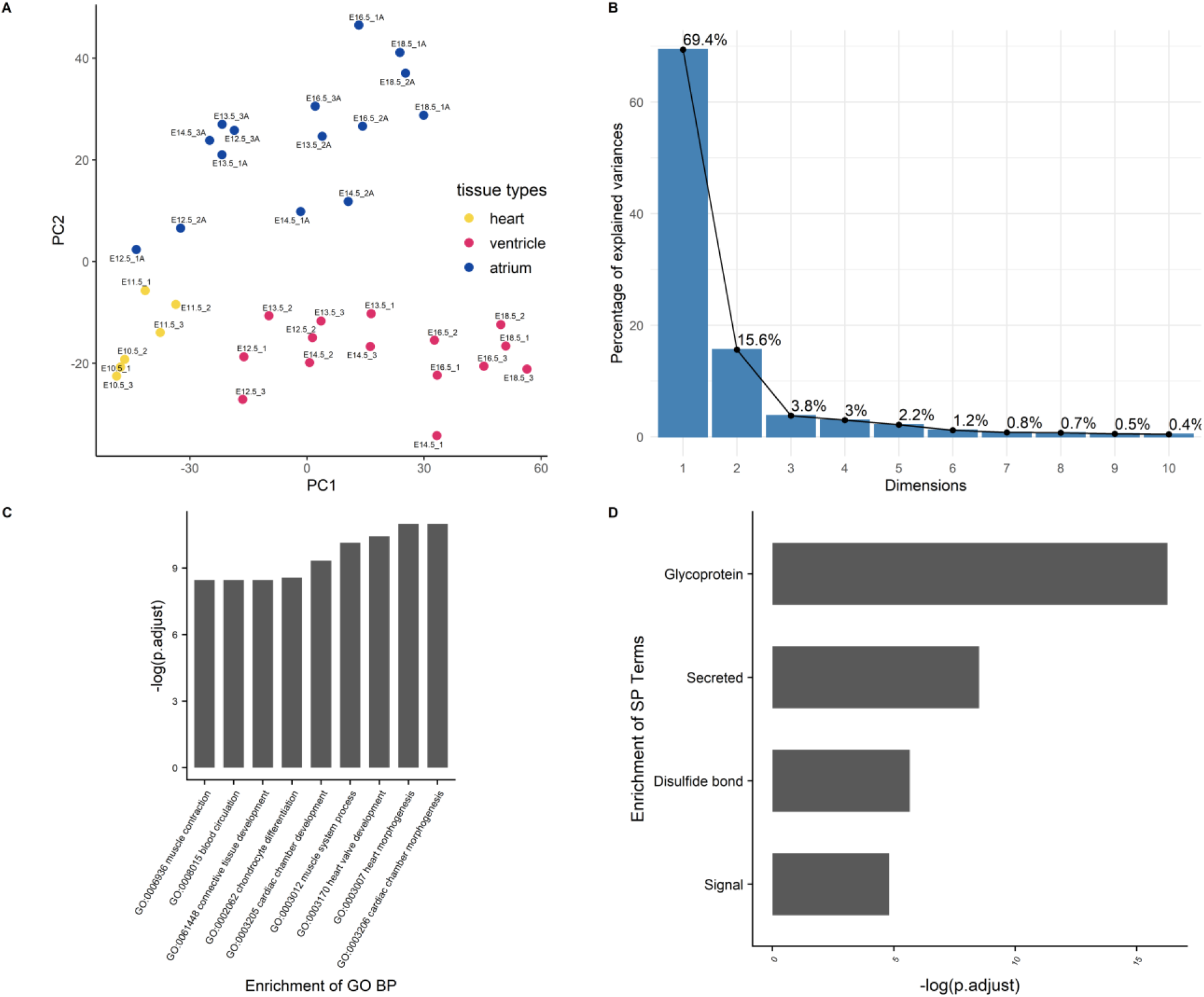
Curation of atrium specific genes from the benchmark set GSE1479. The dataset is a microarray data of high quality generated by CardioGenomics. The data-mining methods were used to find the chamber specific genes from the developmental stages. A) The principle component analysis was performed to check the distribution of the atrium/ventricle groups in the experiment samples. Blue dots were samples from the atrium group. Yellow dots were samples from whole heart group at early stages. Red dots were samples from the ventricle group. These three kinds of tissue sample groups from different stages can be clearly separated by components PC1 and PC2. B) After filtering the chamber specific genes using the differential expression analysis and hierarchical clustering, an PCA procedure using the updated dataset was performed and the corresponding scree plot was generated. The new PC1 can explain nearly 70% variability of the data. C) Atrium genes were analyzed using clusterProfiler and D) DAVID pipeline respectively. The enriched GO Biological Process (BP) annotation and enriched term is in consistent with our expectation, indicating that the chamber gene classification is of high accuracy.

### Expression of atrium gene set was rekindled in myocardial infarction

Data set GSE775 were re-analyzed using above defined set for the atrium genes. This set included spatiotemporal result by the microarray design. The original study included the time-serials samples and the experiment also included three myocardial regions^14^. Reactivation of *natriuretic peptide A* (*Nppa*) and *smooth muscle actin alpha 2 (Acta2*) is the golden standard to assess the quality of myocardial infarction model. The increased expression of both indicated that the mouse model is qualified to downstream analysis (Figure 2A & Figure 2B). Expression of the atrium hallmark gene *Sln* was not statistically significant, however. The manual curated atrium genes plus *Nppa* were plotted using the heatmap. Only heart stress indicator, *Nppa*, was fluctuated with time change. The GSEA analysis using the defined set was then performed to assess the phenotype change. The result showed that the increased expression of atrium genes (Figure 2D) with statistical significance.

**Figure 2.**
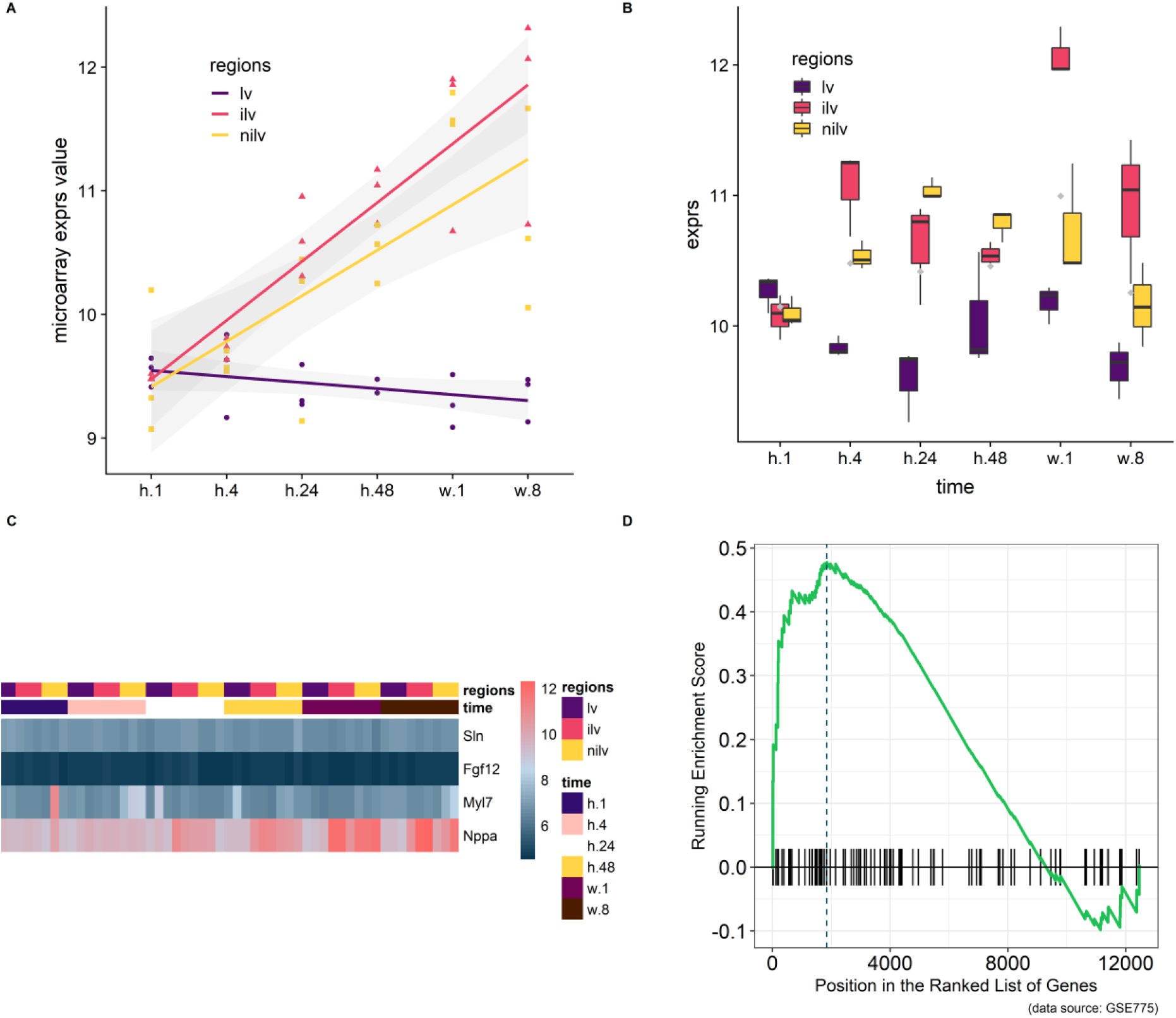
Using atrium gene set in myocardial infarction model (GSE775) and the corresponding GSEA result. A) *Nppa* as the golden heart hypertrophy marker was assayed at the whole time points and regression line was generated to show the upregulation. B) *Acta2* as the alternative marker was used to corroborate the previous finding. C) A small group of manually curated atrium genes was displayed alongside with *Nppa*. The wave-like expression pattern of *Nppa* is in consistent with the previous reports. However, the atrium genes were kept no fluctuate. D) GSEA procedure were performed and result showed that the expression level of the atrium dataset at stage week 1 were increased significantly (*P* = 0.00501).

### Expression of atrium gene set was enlightened in heart regeneration

Data set GSE66403 was used as the biological and technique replicate to infer the phenotype transformation during the heart regeneration. However, the new data set is much more sensitive as the RNA-seq have more accurate measurement. The chamber specific gene set, which was manually curated from the published literature, was used as an independent set to deduce the phenotype change. In the myocardial infarction dataset, the expression of atrium and ventricle genes was not altered. However, the GSEA analysis indicated that the expression of atrium genes are increased, showing that the cell physiological character tend to be atrium cell instead of ventricle cell. While as an independent replicate, the isolated ventricle cells in the heart regeneration dataset showed that the expression of this atrium gene marker was increased and statistically significant. Particularly, expression of the *Sln* was significantly increased (P < 0.05, Figure 3D).

**Figure 3.**
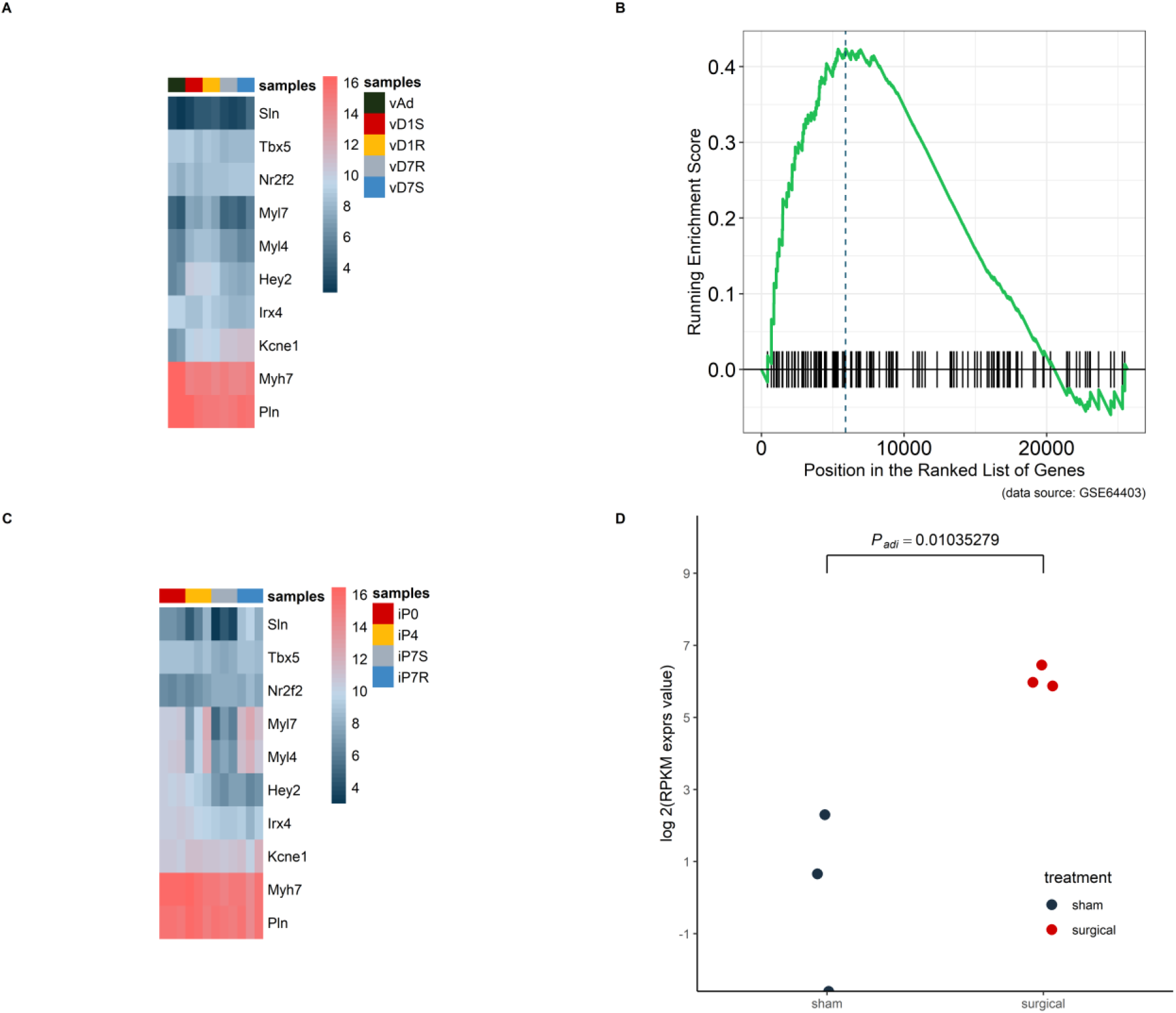
Atrium gene expression pattern were replicated in an independent dataset GSE64403. A) In the heart ventricle tissues, the atrium genes were not significantly increased and the same were ventricle genes. B) The GSEA analysis result showed the increased expression pattern at day-7 with statistical significance (*P* = 0.0262). C) The same group of chamber genes was analyzed in the isolated heart cells, which did not include other lineage cells. The upregulated pattern within the heatmap can be identified in *Sln*, *Myl7* and *Myl4*. D) The golden marker of heart atrium, *Sln* was analyzed at day-7 and the expression level was compared between surgical and sham group. The upregulated *Sln* expression was statistically significant (*P* < 0.05)

### RA signaling upstream of patterning was activated

To trace the upstream event, retinoid acid pathway was explored in both datasets. Retinoic acid (RA) is a vitamin A metabolite that acts as a morphogen during body development. The retinoid pathway was monitored in the present analysis since it plays a role in cardiac chamber patterning. *Aldh1a2* (also named *Radlh2*), component of the RA signaling was activated in the epicardium during zebrafish heart regeneration^23^. *Aldh1a2* was increased in these two datasets. The expression of *Aldh1a2* in the infarction site compared with ventricle sham site or non-injury site was increased significantly at stages h.24 and h.48 respectively (Figure 4A). However, in the regeneration model, the expression of *Aldh1a2* was only significantly increased in the surgery heart compared with the normal heart (Figure 4B).

**Figure 4.**
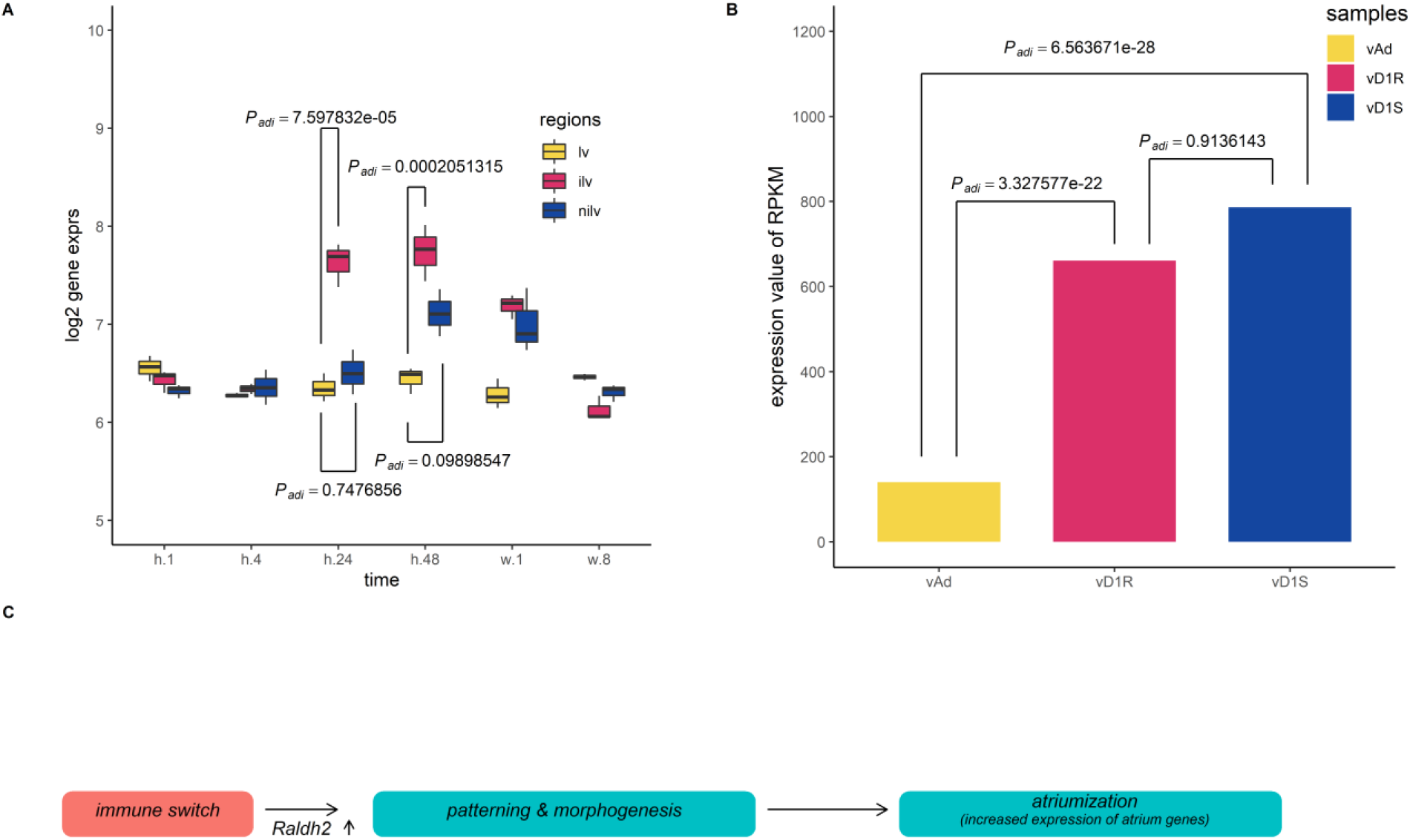
Multiple comparison of *Aldh1a2* expression between different groups in dataset GSE64403. The *P* values were adjusted using the nbinomTest method in DESeq2 package. A) *X*-axis stands for the sampling stages at 1 hour, 8 hours, 24 hours, 48 hours, 1 week and 8 weeks in an operated mouse. Lv stands for left ventricle. Ilv stands for the infarcted and nilv is taken from non-infarcted left ventricle. B) vAd was the mouse sample from the region of adult ventricular myocardium. vD7S was the mouse sample from the region of 7 days post-sham surgery ventricular myocardium. vD7R was the sample from the region of 7 days post-sham surgery ventricular myocardium. Expression of *Aldh1a2* from vD7R or vD7S was increased compared with that from vAd (*P_adj_* < 0.001). C) The graph summarized the inference of the present results, which indicated that the initial immune response stimulates the development patterning events during heart injury.

## Discussions

We rigorously defined the atrium gene sets, including ventricle gene set using a benchmark dataset GSE1479. The myocardial infarction (GSE775) and heart regeneration (GSE64403) datasets were explored using data-mining methods. The results showed that the expression of atrium genes was increased in the harvested heart ventricle tissue. This observation can be explained by three interpretations. One possible mechanism is ventricular-to-atrial *trans*-differentiation, leading to the local increase of atrium cells. The second is atrium-like progenitor cell proliferation. The third may be the atrium cell migration from heart atrium. We cannot exclude the other possibility. The discovery of this phenomena is solely dependent on the re-analysis of these public data.

Verification of chamber specific cell requires three pieces of evidence in wet procedure. They include gene expression pattern, electrophysiological properties and calcium sparks^24,25^. In the present study, a predefined atrium set was used in GSEA analysis and *in facto* atrium cell marker, *Sln*, was used as a second evidence. Adolfo J. de Bold proposed that, based upon the association of *Nppa* with the atrial-specific granules, the concept of the heart as an endocrine organ^26^. We also found in the present unbiased bioinformatics approach that atrial cardiomyocytes displayed secretory phenotype. The heart is regarded as a endocrine organ thirty years ago, and secretome profiling also the hallmark during cardiomyogenesis^27^. Expression of *Nppa* was an early and specific marker for the differentiating working myocardium of the atria and ventricles of the developing heart^28^. However, our statistical analysis did not support *Nppa* as the hallmark of atrium phenotype. Instead, *Nppa* is a true marker in clinical investigation of cardiovascular diseases. Two separated enhancers which govern independent response during development and disease^28^. Instead, recent study revealed that *Sln* is the true lineage marker of atrium cells and have a conserved expression pattern. Early work showed that *Sln* was up-regulated during development and down-regulated by cardiac hypertrophy^20^. Thus, expression of *Sln* is a perfect inner control in our study. We found that the expression of *Sln* mRNA was detected selectively in the atria and not in myocardial infarction.

We hypothesized that the expression of atrium genes would be increased in response to the cardiac damage and myocardial infarction (MI) is permanent damage to the heart muscle. The microarray data, GSE775, was recruited to support our assumption. This mouse dataset is a time series intended to compare normal functioning left ventricles with infarcted (ilv) and non-infarcted left ventricles (nilv). Ilv samples are taken from the region between the LAD artery and the apex on a mouse with myocardial infarction. Nilv samples are taken from the region above the infarction and the left ventricle (lv) samples mimic that region in a sham mouse. To double check the quality of the dataset and analysis pipeline, the *Nppa* expression was assayed as it is the golden standard of cardiac diseases. Reactivation of ventricular *Nppa* expression is part of a conserved adaptive change in molecular phenotype in response to heart failure that serves both diagnostic and potential therapeutic options^29,30^. The expression levels of *Nppa* was regressed across the whole time points and upregulated in the ilv or nilv samples while that in lv sample (sham counterpart) was kept same.

To further confirm the finding that the myocardial infarction model was successful, the expression of *Acta2* was checked as it is an alternative marker which was used to corroborate the finding^31^. The upregulation of both *Nppa* and *Acta2* at early stages indicates that the mouse model truly exhibits the disease progression. To test expression level of the atrium genes, *Nppa* and other manually curated atrium genes^20,32,33^ were plotted in the same heatmap. The wave-like expression pattern of *Nppa* (Figure 2C) is in consistent with the previous analysis (Figure 2A). However, the atrium genes were kept no change. The *Sln* expression is consistent with the previous finding that the expression in the heart ventricle is not upregulated during cardiac hypertrophic remodeling. GSEA analysis was then undertaken to increase the power the detect the expression direction. The result showed that the expression levels of the atrium dataset were increased significantly.

To further confirm our finding that expression of atrium gene set was upregulated, atrium gene expression pattern was verified in an independent dataset GSE64403, which utilized deep-sequencing method and heart regeneration model. In the heart ventricle tissues, the atrium genes were not scientifically increased and the same were ventricle genes (Figure 3A). GSEA analysis was conducted and showed that the increased expression of atrium genes is of above statistical significance. However, the harvested ventricle tissues included other cell lineages besides cardiac ventricle cells. To exclude this influence and identify the true source, the same group of chamber genes was analyzed in the isolated heart ventricle cells. The increased expression pattern can be identified in the authentic atrium markers, *Sln*, *Myl7* and *Myl4* (Figure 3C). The golden marker of atrium gene, *Sln* was analysis and compared between surgical and sham group at day 1. The upregulated *Sln* expression was statistically significant (*P* < 0.05). In the present study, we examined the effects of development and cardiac hypertrophy on the expression of *Sln* mRNA in heart. We found that the expression of *Sln* was detected selectively in the atria. Furthermore, *Sln* was upregulated during development and down-regulated by cardiac hypertrophy^20^.

To ask what the upstream event of atrium gene reactivation, multiple comparisons of *Aldh1a2* expression between different groups in data set GSE64403 were performed. Early study in zebrafish revealed that the injured heart at the epicardium exhibited increased expression of *Radhl2* using mRNA probe hybridization. The *P* values were adjusted using the nbinomTest method in DESeq2 package. The vAd sample was from the region of adult mouse ventricular myocardium. The vD7S was from the region of 7 days post-sham surgery ventricular myocardium. The vD7R was from the region of 7 days post-sham surgery ventricular myocardium. Expression of *Aldh1a2* from vD7R or vD7S was increased compared with that from vAd (*Padj* < 0.001). *Hox* genes and retinoid receptors were also checked and they were at different stages upregulated. These results indicate that at earliest stage of myocardial infarction or heart regeneration, immune response elicited the patterning process to be reactivated through retinoid acid signaling pathway, which mirrors the developmental progress in cardiac development.

Numerous studies propose that the heart regeneration involved in re-activation of the development pathways^9^. But this conclusion is not based upon the quantitative data. We here used the bioinformatic analysis to infer that the ventricle-atrium *trans*-differentiation might occur during the heart regeneration or myocardial infarction in mouse model due to the activation of body patterning event.

## Conclusions

The present approach using the public datasets and data-mining methods indicated that the increased expression of atrium specific genes is the hallmark of local response to the heart injures. The so-called atriumization then revealed possible phenotype transformation during heart regeneration. This kind of gene expression upregulation suggested that the heart ventricle cells preferred to withdrawing their default physiological character and make an attempt by the outside morphogen signal to turn into atrium-like cells. Here, the regression to the atrium cell might be the first step toward to local immune response and possible the atavism that atrium cardiac chamber is the default evolutionary appearance order^34^. The cell-level reprogramming might be the fundamental mechanism of priming heart regeneration.

Impediment of the process probably resulted in cardiomyocyte loss, leading to myocardial infarction or regeneration failure. Our work is only based upon the public data and needs to be addressed by wet verification. The preliminary results will shed light on the future investigation on regeneration medicine, however.

## Acknowledgments

This manuscript has been released as a pre-print at bioRxiv^35^ (Yisong Zhen)

